# Heteromeric amyloid filaments of ANXA11 and TDP-43 in FTLD-TDP Type C

**DOI:** 10.1101/2024.06.25.600403

**Authors:** Diana Arseni, Takashi Nonaka, Max H. Jacobsen, Alexey G. Murzin, Laura Cracco, Sew Y. Peak-Chew, Holly J. Garringer, Ito Kawakami, Hisaomi Suzuki, Misumoto Onaya, Yuko Saito, Shigeo Murayama, Changiz Geula, Ruben Vidal, Kathy L. Newell, Marsel Mesulam, Bernardino Ghetti, Masato Hasegawa, Benjamin Ryskeldi-Falcon

## Abstract

Neurodegenerative diseases are characterised by the abnormal filamentous assembly of specific proteins in the central nervous system^1^. Human genetic studies established a causal role for protein assembly in neurodegeneration^2^. However, the underlying molecular mechanisms remain largely unknown, which is limiting progress in developing clinical tools for these diseases. Recent advances in electron cryo-microscopy (cryo-EM) have enabled the structures of the protein filaments to be determined from patient brains^1^. All diseases studied to date have been characterised by the self-assembly of a single intracellular protein in homomeric amyloid filaments, including that of TAR DNA-binding protein 43 (TDP-43) in amyotrophic lateral sclerosis (ALS) and frontotemporal lobar degeneration with TDP-43 inclusions (FTLD-TDP) Types A and B^3,4^. Here, we used cryo-EM to determine filament structures from the brains of individuals with FTLD-TDP Type C, one of the most common forms of sporadic FTLD-TDP. Unexpectedly, the structures revealed that a second protein, annexin A11 (ANXA11), co-assembles with TDP-43 in heteromeric amyloid filaments. The ordered filament fold is formed by TDP-43 residues G282/284–N345 and ANXA11 residues L39–L74 from their respective low-complexity domains (LCDs). Regions of TDP-43 and ANXA11 previously implicated in protein-protein interactions form an extensive hydrophobic interface at the centre of the filament fold. Immunoblots of the filaments revealed that the majority of ANXA11 exists as a ∼22 kDa N-terminal fragment (NTF) lacking the annexin core domain. Immunohistochemistry of brain sections confirmed the co-localisation of ANXA11 and TDP-43 in inclusions, redefining the histopathology of FTLD-TDP Type C. This work establishes a central role for ANXA11 in FTLD-TDP Type C. The unprecedented formation of heteromeric amyloid filaments in human brain revises our understanding of amyloid assembly and may be of significance for the pathogenesis of neurodegenerative diseases.

## MAIN

The abnormal assembly of TDP-43 is the hallmark of multiple neurodegenerative diseases, including FTLD-TDP, ALS and limbic predominant age-related TDP-43 encephalopathy (LATE), as well as of inclusion body myopathy^5–8^. Currently, there are no effective means to diagnose or treat these diseases. While the wild-type protein assembles in the majority of disease cases, pathogenic variants in the gene encoding TDP-43, *TARDBP*, that increase the propensity of the mutated protein to assemble indicate a causal role for assembly in disease^9,10^.

Four types of FTLD-TDP, designated A–D, are distinguished by the distribution of assembled TDP-43 in the brain and are associated with different frontotemporal dementias (FTD)^11^. In FTLD-TDP Type C, neocortical assembled TDP-43 is predominantly distributed in elongated inclusions within dystrophic neurites (DNs), which are somewhat more abundant in the superficial cortical layers^11^. This contrasts with the other three types of FTLD-TDP, where assembled TDP-43 is mainly found in inclusions within neuronal soma. Compact neuronal cytoplasmic inclusions (NCIs) of assembled TDP-43 are also present in the hippocampal dentate gyrus and striatum in FTLD-TDP Type C. FTLD-TDP Type C is most-frequently associated with semantic variant primary progressive aphasia (svPPA), which is characterised by selective neurodegeneration of the anterior temporal lobes and word comprehension impairments^12^. Unlike the other types of FTLD, for which pathogenic genetic variants account for a large proportion of cases, no such variation has been associated with FTLD-TDP Type C, which limits our understanding of its pathogenesis.

In its native form, TDP-43 is a ubiquitous RNA-binding protein with multiple regulatory roles in RNA processing^13^. It predominantly localises to ribonucleoprotein (RNP) granules in the nucleus, but also undergoes nucleocytoplasmic shuttling and can be found in cytoplasmic RNP granules^14^. TDP-43 is comprised of an N-terminal dishevelled and axin (DIX) domain, a nuclear localisation signal, tandem RNA recognition motifs (RRMs) and a C-terminal LCD. The latter contains three distinct regions enriched in glycine, hydrophobic residues, and glutamine and asparagine (Q/N-rich region).

In disease, assembled TDP-43 comprises the full-length protein and truncated C-terminal fragments (CTFs), both of which are abnormally ubiquitylated and phosphorylated^5,6^. The assemblies are filamentous^15–19^, but bind amyloidophilic dyes such as Thioflavin S poorly^20^. Using cryo-EM, we previously established that TDP-43 filaments in ALS and FTLD-TDP Types A and B are amyloids^3,4^.

Amyloid filaments are characterised by highly-stable ordered folds containing parallel, in-register intermolecular β-sheets in line with the filament axis^21^. They have been defined as self-assemblies of identical or near-identical protein sequences. *In vitro* studies have shown that a given protein sequence has the potential to form a vast number of different filament folds^22^. Remarkably, each disease studied by cryo-EM to date has been characterised by specific filament folds^1^. This suggests that distinct processes lead to the formation of disease-characteristic filament folds in patient brain.

The ordered folds of TDP-43 filaments in ALS and FTLD-TDP Types A and B are comprised entirely of the N-terminal half of the TDP-43 LCD, with flanking sequences forming a fuzzy coat. Despite this, they are folded differently between FTLD-TDP Type A and the disease continuum of ALS and FTLD-TDP Type B. The structures of TDP-43 filaments in other diseases are not known. Here, we used cryo-EM to determine the structures of filaments from the brains of individuals with FTLD-TDP Type C.

### Extraction of filaments from FTLD-TDP Type C

We extracted filaments from the prefrontal and temporal cortices of four individuals with FTLD-TDP Type C (Extended Data Table 1), according to the method we previously used for ALS and FTLD-TDP Types A and B^3,4^. Immunohistochemistry of prefrontal cortex brain sections from these individuals confirmed the presence of abundant elongated TDP-43-inclusions within DNs, in the absence of abundant inclusions in neuronal soma (Extended Data Fig. 1a), diagnostic of FTLD-TDP Type C^11^. Immuno-gold negative-stain electron microscopy (immuno-EM) of the extracts confirmed the presence of TDP-43-immunoreactive filaments (Extended Data Fig. 1b), as previously reported in extracts^23^, and observed *in situ* within DNs of individuals with FTLD-TDP Type C^16,17^. These filaments were readily identifiable in cryo-EM images of the extracts (Fig. 1a).

**Fig. 1:**
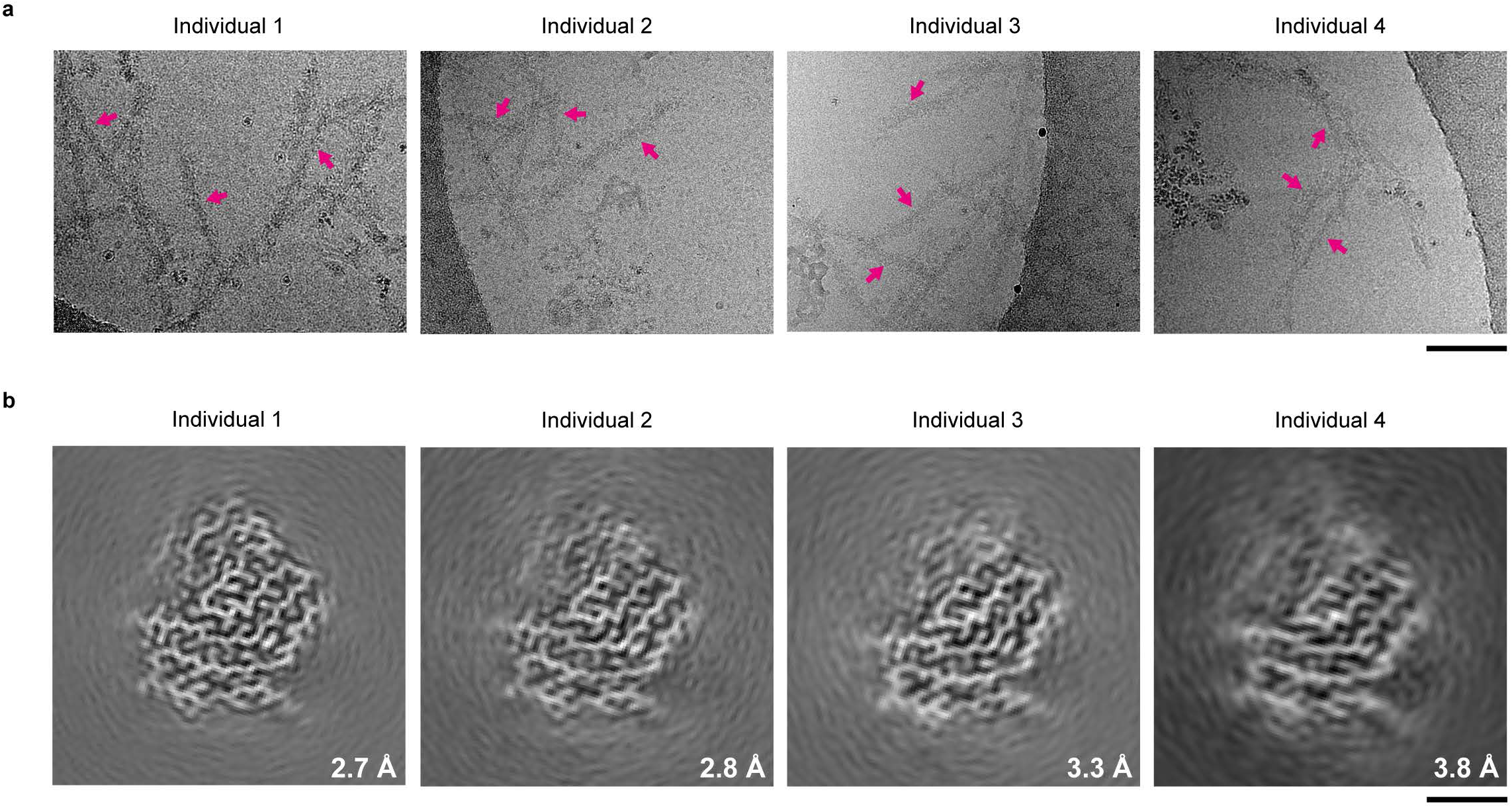
Cryo-EM of filaments from individuals with FTLD-TDP Type C. **a**, Representative cryo-EM images of filaments extracted from the prefrontal and temporal cortex of four individuals with FTLD-TDP Type C. Examples of filaments are indicated with arrows. The filaments were identified by their width of ∼15 nm, helical crossover distance of ∼50 nm and granular surfaces. Scale bar, 100 nm. **b,** Cryo-EM reconstructions of filaments from the four individuals with FTLD-TDP Type C, shown as central slices perpendicular to the helical axis. All four reconstructions have the same filament fold. The resolution of each reconstruction is indicated. Scale bar, 2 nm.

In addition to TDP-43 filaments, we observed occasional amyloid-β (Aβ) filaments in the cryo-EM images for all individuals and tau paired helical filaments for individuals 1–3, which were evident based on their distinct widths and helical crossover distances (Extended Data Fig. 2). This is consistent with the presence of sparse Aβ plaques and tau tangles in these individuals (Extended Data Table 1). We also observed distinct transmembrane protein 106B (TMEM106B) filaments for individuals 1–3, but not 4 (Extended Data Fig. 2). This is consistent with the previously-reported age-dependent accumulation of TMEM106B filaments in human brain^24^. Individuals 1–3 were all ≥74 years-old, whereas individual 4 died at 59 years-of-age (Extended Data Table 1). The presence of these filaments in the extracts was supported by mass spectrometry, which identified peptides corresponding to their ordered folds (Supplementary Table 1).

### Cryo-EM reveals that ANXA11 forms heteromeric amyloid filaments with TDP-43

We collected 296,660 cryo-EM images of the filament extracts and used helical reconstruction to determine the structures of the TDP-43-immunoreactive filaments from each of the four individuals independently, achieving resolutions of up to 2.7 Å (Fig. 1b, Extended Data Fig. 3 and Extended Data Table 2). We found a single filament type with an identical ordered fold among the four individuals, suggesting that this filament fold characterises FTLD-TDP Type C. The fold comprises two discrete complementary protein chains of different length and conformation (Fig 2a and b). Three-dimensional classification of the largest dataset of cryo-EM filament segments, for individual 1, revealed a variable region at the distal end of the longer chain that adopts two alternative conformations (Extended Data Fig. 3 and Extended Data Table 2). This fold, which resembles a kite (deltoid) in profile, is distinct from the double-spiral fold of ALS and FTLD-TDP Type B and the chevron fold of FTLD-TDP Type A (Extended Data Fig. 4a)^3,4^.

**Fig. 2:**
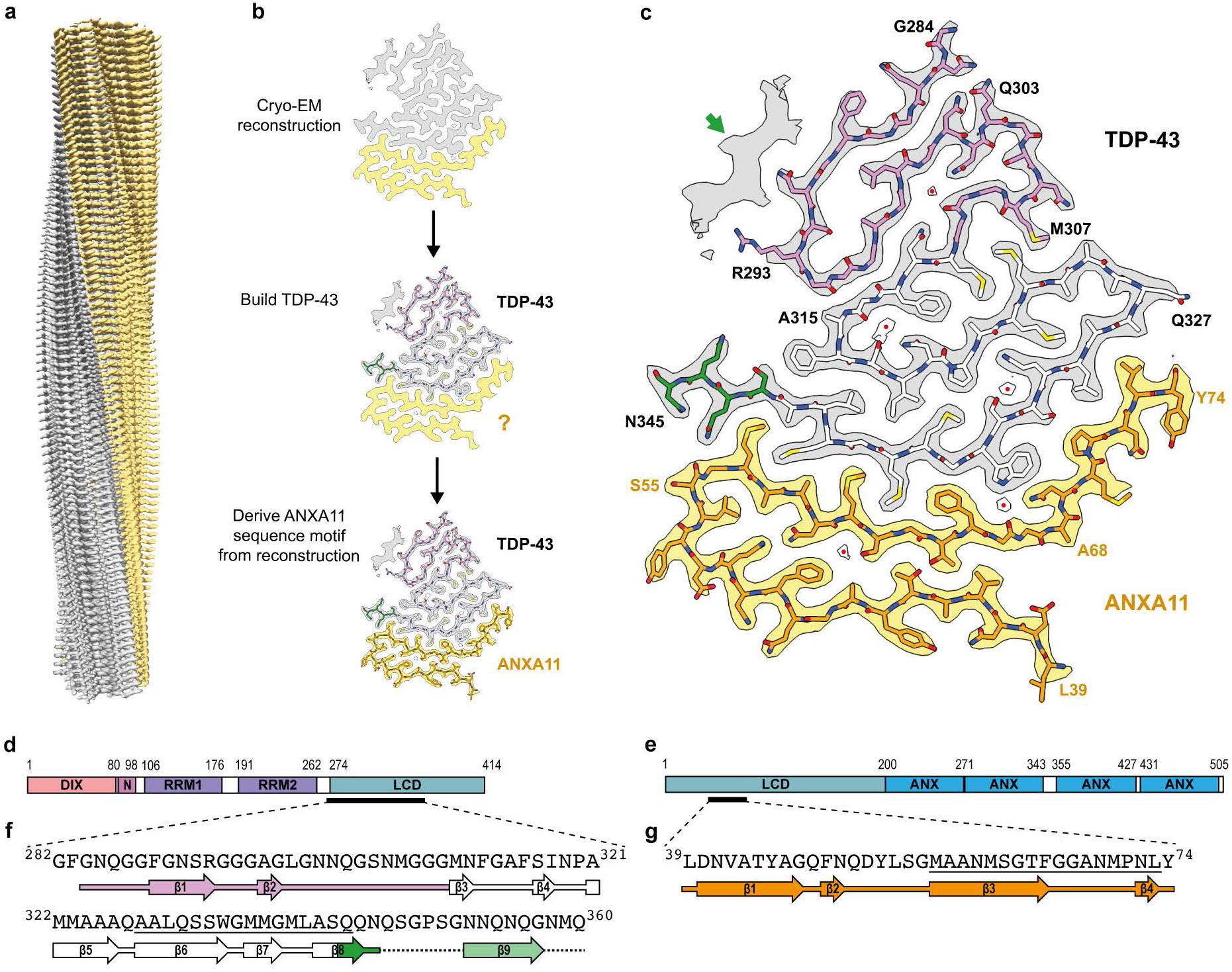
Cryo-EM structure of heteromeric amyloid filaments of ANXA11 and TDP-43 from FTLD-TDP Type C. **a**, Cryo-EM reconstruction of the left-handed filaments of ANXA11 and TDP-43 from FTLD-TDP Type C, shown parallel to the helical axis. **b,** Identification of TDP-43 and ANXA11 chains in the ordered filament fold. ANXA11 was identified by deriving a sequence motif directly from well-resolved amino acid side chain densities in the cryo-EM reconstruction (see Methods). **c,** Cryo-EM reconstruction and atomic model of the filaments, shown for single TDP-43 and ANXA11 chains perpendicular to the helical axis. The green arrow indicates an isolated peptide consistent with TDP-43 residues N352-G357. Buried ordered solvent is indicated with red dots. **d and e,** Domain organisation of TDP-43 **(d)** and ANXA11 **(e).** DIX, dishevelled and axin domain; **N,** nuclear localisation signal; RRM, RNA-recognition motif; LCD, low-complexity domain; ANX, annexin repeat. The black lines indicate the regions that form the filament fold. **f and g,** Amino acid sequence alignment of the secondary structure elements of the TDP-43 **(f)** and ANXA11 **(g)** chains. Arrows indicate -strands. The sequences that form the interface between TDP-43 and ANXA11 are underlined. **a-c,** Cryo-EM density for TDP-43 is in grey and ANXA11 is in yellow. **b, c, f and g,** The TDP-43 glycine-rich (G284-G310, magenta), hydrophobic (M311-S342, white) and QIN-rich (Q343-Q345, green) regions are highlighted. ANXA11 is shown in orange.

The protein backbone and amino acid side chains were unambiguously resolved in our 2.7 Å cryo-EM reconstruction, thereby enabling us to build an accurate atomic model of the filament fold (Fig. 2a–c, Extended Data Fig. 3 and Extended Data Table 2). The filaments have a left-handed helical twist, in contrast with the right-handed filaments of ALS and FTLD-TDP Types A and B^3,4^. The longer chain comprises the TDP-43 sequence G282/284–N345 from its LCD (Fig. 2b–d and f). This is similar to the sequences that form the TDP-43 filament folds of ALS and FTLD-TDP Types A and B, but lacks 15 residues from the Q/N-rich region (Q346–Q360) (Extended Data Fig. 4b).

Unexpectedly, the shorter second chain could not accommodate any sequences from TDP-43. To identify the protein sequences forming this chain, we derived a sequence motif directly from the well-resolved amino acid side chain densities of the cryo-EM reconstructions, which returned a single hit when searched against reference proteomes (Fig. 2b and Methods). This revealed that the chain belongs to the calcium-dependent phospholipid-binding protein ANXA11 (ref. ^25^), comprising the sequence L39–L74 from its N-terminal LCD (Fig. 2b, c, e and g). Among the 12 human annexins, ANXA11 is unique in possessing an LCD^26^. It is enriched in proline residues, apart from the region spanning I37–M70, which lacks proline. The ANXA11 chain comprises almost all of this region, together with four additional residues toward the C-terminus. As such, there is only a single proline (P71) in its sequence. We confirmed the presence of both ANXA11 and TDP-43 in the filaments of FTLD-TDP Type C using double-labelling immuno-EM (Extended Data Fig. 5). Furthermore, mass spectrometry of the filament extracts detected ANXA11, in addition to TDP-43, with an enrichment of peptides from the ordered filament fold (Supplementary Table 1). All previous structures of neurodegenerative disease-associated assemblies from patient brain have been self-assemblies of a single protein within homomeric amyloid filaments^1^. This is the first evidence that amyloid filaments can be heteromeric in human brain. This work establishes an unprecedented central role for ANXA11 in FTLD-TDP Type C.

#### The heteromeric filament fold of ANXA11 and TDP-43

One ANXA11 chain and one TDP-43 chain complement each other in the ordered fold of the heteromeric amyloid filaments of FTLD-TDP Type C (Figure 2c and Extended Data Fig. 6a). Residues S55–Y74 from ANXA11 and Q327–N345 from TDP-43 form the most striking interface of the filament fold. This interface comprises two antiparallel layers that associate tightly over their entire lengths, bending in the middle toward the TDP-43 layer (Fig. 3a). Of the 15 amino acid side chains that participate in this interface, 12 (80%) are hydrophobic (Fig. 3b). At one end of the interface, the side chain of TDP-43 residue Q343 hydrogen bonds to the main chain of ANXA11 residue S55 (Fig. 3c). In the middle of the interface, there is an ordered solvent molecule that mediates interactions between the polar groups of TDP-43 residue W334 and ANXA11 residues G66 and N69 (Fig. 3a).

**Fig. 3:**
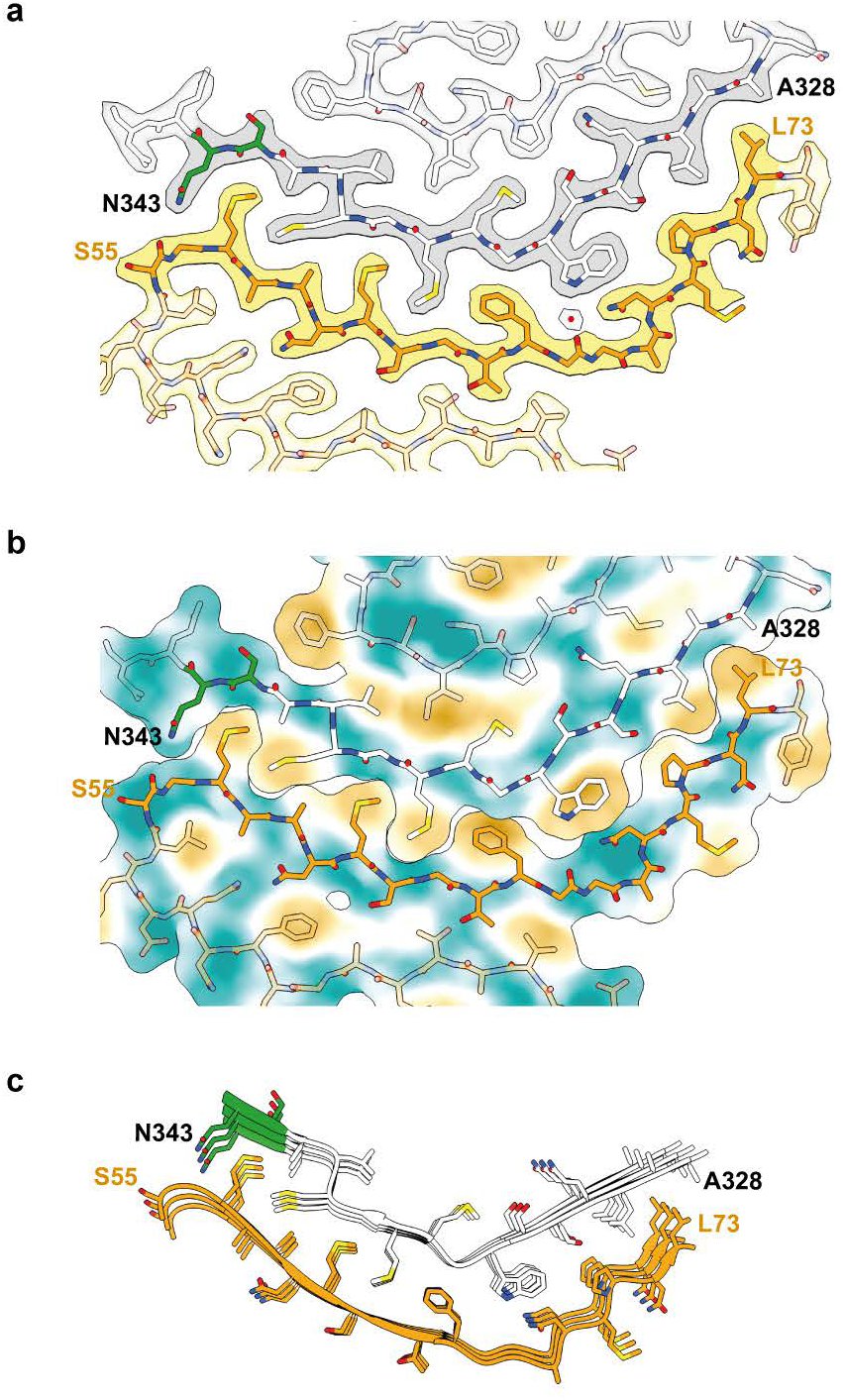
ANXA11 and TDP-43 interface in heteromeric amyloid filaments from FTLD-TDP Type C. **a and b**, Overlay of the cryo-EM reconstruction **(a)** and hydrophobicity surface plot (yellow, most hydrophobic; teal, least hydrophobic) **(b)** with the atomic model of the filaments, focussed on the interface between ANXA11 and TDP-43 and shown for single ANXA11 and TDP-43 chains perpendicular to the helical axis. Cryo-EM density for TDP-43 is in grey and ANXA11 is in yellow. Buried ordered solvent is indicated with a red dot. **c,** Atomic model of the interface between ANXA11 and TDP-43, shown for three molecular layers perpedicular to the helical axis.

The remaining ANXA11 residues, L39–L54, fold back against the opposite side of the ANXA11 interface layer, forming a second layer that extends up to the bend in the interface (Fig. 2c and Extended Data Fig. 6a). The two ANXA11 layers associate through a mixture of non-polar and polar interactions, including sparse hydrogen bonds between Q51 and N60, and between the two threonine residues (T44 and T64) (Extended Data Fig. 6b).

The remaining TDP-43 residues, G282/284–A326, fold inside the bend of the TDP-43 interface layer in a serpentine arrangement (Fig. 2c and Extended Data Fig. 6a). The rest of the TDP-43 hydrophobic region (M311–A326) forms a second layer and half of a third layer. These layers associate through hydrophobic interactions as well as through sparse hydrogen bonding between neutral polar residues (Extended Data Fig. 6b and c). The TDP-43 glycine-rich region (G282/284–G310) completes the third layer and adds two more layers in the main conformation of the fold and one pleated layer in the alternative conformation (Extended Data Fig. 7a–c). Similar structural variation of the glycine-rich region was previously observed in the TDP-43 filament fold of FTLD-TDP Type A^4^.

In the alternative conformation, residues G283–R293 form a compact substructure identical to that of the TDP-43 filament fold of FTLD-TDP Type A^4^ (Extended Data Fig. 7d). In the latter, the side chain of R293 is buried without charge compensation, whereas in the structure reported here ordered solvent molecules are present adjacent to the R293 guanidino group, possibly acting as counterions (Extended Data Fig. 7a). As in FTLD-TDP Type A, we found citrullination of R293 using mass spectrometry of the extracted filaments (Supplementary Table 1), which would facilitate the formation of this compact substructure by removing the charge of R293. This suggests that citrullination of R293 may be of broad significance for TDP-43 assembly in disease.

In the main conformation, we observed an isolated density island that is likely to correspond to a peptide of approximately six residues engaging in zipper packing with the β-strand formed by F289–R293 in the TDP-43 glycine-rich region (Fig. 2c). This isolated density is consistent with TDP-43 residues N352–G357 and likely represents an extension from N345 at the C-terminus of the fold, with the intervening residues (Q346–G351) being disordered (Fig. 2f). This density island was absent in the alternative conformation of the glycine-rich region (Extended Data Fig. 7a).

Both the TDP-43 and ANXA11 chains of the heteromeric filaments are stabilised along the helical axis by hydrogen bonding within intermolecular parallel in-register β-sheets and glutamine/asparagine side chain ladders, as well as staggered stacking interactions of aromatic side chains, characteristic of amyloid filaments (Extended Data Fig. 6b and d).

### C-terminal truncation of ANXA11 in the heteromeric filaments of FTLD-TDP Type C

Abnormal CTFs of TDP-43, in addition to the full-length protein, are present in disease assemblies^5,6^. To identify the molecular species of ANXA11 in the heteromeric filaments of FTLD-TDP Type C, we performed immunoblot analysis of filament extracts (Fig. 4a). This

**Fig. 4:**
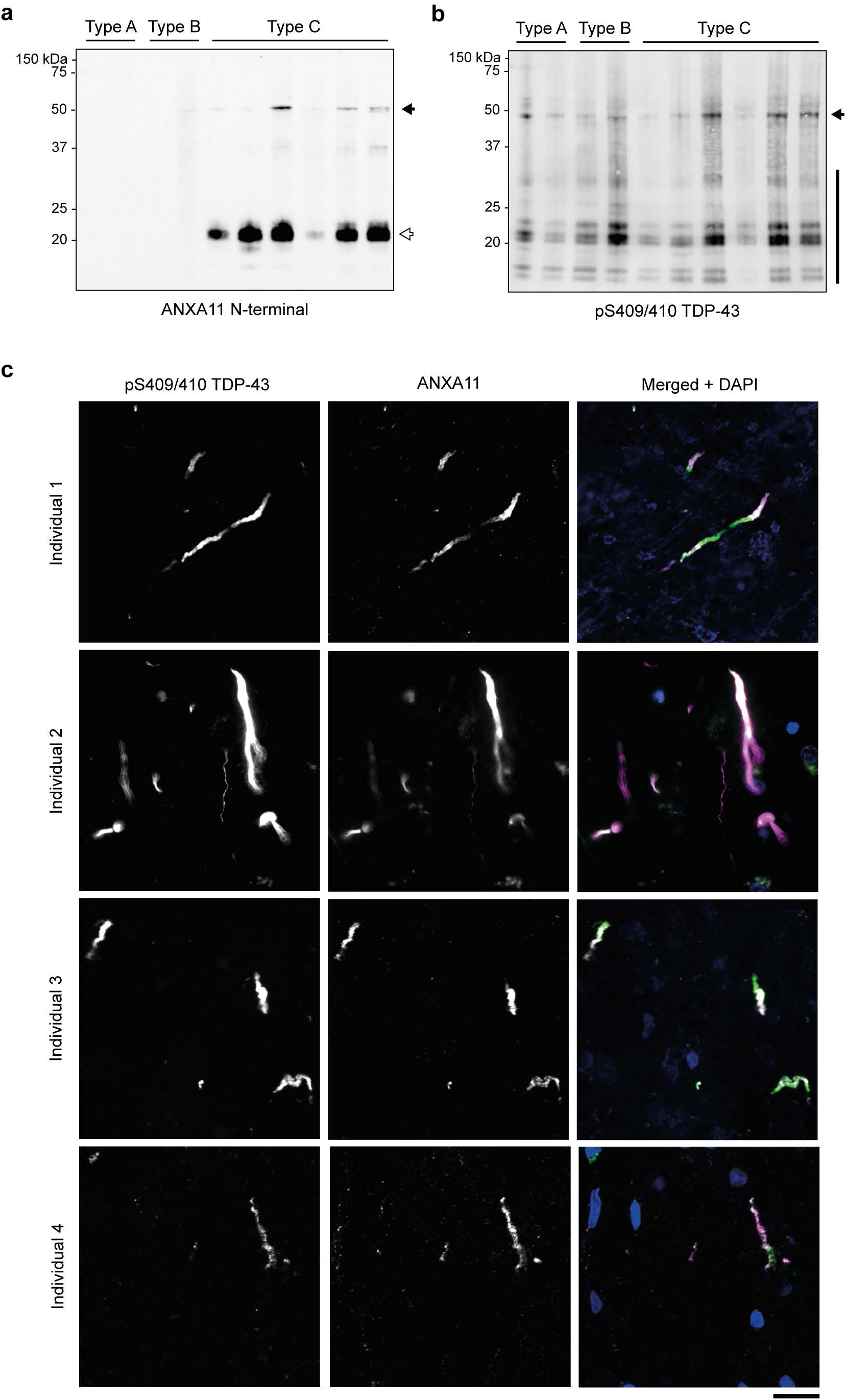
Molecular pathology of ANXA11 in FTLD-TDP Type C. **a and b**, lmmunoblot analysis of filament extracts from the prefrontal cortex of individuals with FTLD-TDP Types A (2), B (2) and C (6) using antibodies against N-terminal ANXA11 (residues 1-180) **(a)** and pS409/410 TDP-43 **(b).** A ∼22 kDa ANXA11 NTF (white arrow), as well as a minor population of full length ANXA11 (black arrow), are observed for all individuals with FTLD-TDP Type C, but not for individuals with FTLD-TDP Types A and B. Full length TDP-43 (black arrow) and TDP-43 CTFs (black line) are observed for all individuals. **c,** lmmunohistochemical analysis of prefrontal cortex sections from four individuals with FTLD-TDP Type C using antibodies against pS409/410 TDP-43 and N-terminal ANXA11 (residues 1-180). Individual images for TDP-43 and ANXA11 are shown in greyscale to facilitate comparison, in addition to a merged image showing TDP-43 (green), ANXA11 (magenta) and DAPI (blue) staining. ANXA11 and TDP-43 co-localise with inclusions. Additional immunohistochemical analysis is shown in Extended Data Fig. 8 and 9. Scale bar, 20 µm.

revealed that ANXA11 predominantly existed as an NTF that migrated at ∼22 kDa, in addition to a minor population of full-length (56 kDa) ANXA11 (Fig. 4a). The ANXA11 NTF was not detected on immunoblots of filament extracts from individuals with FTLD-TDP Types A and B (Fig. 4a), in support of a lack of ANXA11 in the homomeric TDP-43 filament folds of these diseases^3,4^. A faint band for full-length ANXA11 was detected for one Type B case with the highest protein load. Full-length TDP-43 and CTFs of ∼18–35 kDa phosphorylated at S409 and S410 were detected for all individuals (Fig 4b), as previously observed^5,6^. These molecular species of ANXA11 and TDP-43 are larger than their sequences in the ordered filament fold (36 residues, ∼3.8 kDa, and 64 residues, ∼6.2 kDa, respectively), indicating that extensive flanking regions extend from the fold to form a fuzzy coat^1^. The ANXA11 NTF has not been reported before and constitutes a new pathological hallmark of FTLD-TDP Type C.

### A redefined histopathology of FTLD-TDP Type C

Owing to our finding of heteromeric filaments composed of both ANXA11 and TDP-43 in FTLD-TDP Type C, we predicted that the TDP-43-immunoreactive inclusions of this disease should also be immunoreactive against ANXA11. Indeed, multiplexed fluorescence immunohistochemistry of prefrontal cortex brain sections from individuals with FTLD-TDP Type C showed co-localisation between the two proteins in DN inclusions (Fig. 4c and Extended Data Fig. 8a). We did not observe co-localisation between ANXA11 and TDP-43-immunoreactive inclusions in prefrontal cortex brain sections from individuals with FTLD-TDP Types A and B (Extended Data Fig. 8b and c), in agreement with a lack of ANXA11 in their homotypic TDP-43 filament structures^3,4^. We also observed co-localisation between TDP-43 and ANXA11 in the NCIs of the hippocampal dentate gyrus of individuals with FTLD-TDP Type C (Extended Data Fig. 9), suggesting that heteromeric amyloid filaments of ANXA11 and TDP-43 are also present in these inclusions. We did not observe inclusions that were immunoreactive against only ANXA11 or only TDP-43, in agreement with our cryo-EM structures in which both proteins co-assemble in heteromeric filaments. ANXA11 immunoreactivity of TDP-43 inclusions redefines the histopathology of FTLD-TDP Type C.

## DISCUSSION

Specific proteins that assemble into filaments in neurodegenerative diseases, including TDP-43, tau and α-synuclein, have been previously identified using biochemical fractionation of inclusions from patient brains^5,6,27–29^, of which they are the major component. Antibodies raised against these proteins have subsequently been used to identify associated diseases. Recent breakthroughs in cryo-EM have enabled the structures of these protein filaments to be determined from patient brains^1^. An additional strength of high-resolution cryo-EM is that it can identify novel protein assemblies, as shown for those of TMEM106B in aging and TAF15 in FTLD^24,30^. Here, using cryo-EM, we unexpectedly discovered that a second protein, ANXA11, co-assembles with TDP-43 heteromeric amyloid filaments in FTLD-TDP Type C. This reveals a central role for ANXA11 in FTLD-TDP Type C. Amyloid filaments were previously defined as self-assemblies of identical or near-identical protein sequences. Indeed, all previously determined structures of neurodegenerative disease-associated assemblies from patient brain have been homomeric amyloid filaments of a single specific protein^1^. Our discovery of heteromeric amyloid filaments composed of ANXA11 and TDP-43 revises this definition and raises several mechanistic hypotheses.

Pathogenic variants in *ANXA11* have been linked to ALS, inclusion body myopathy and FTD^31–34^. This includes svPPA^35^, which is most commonly associated with FTLD-TDP Type C^11^. Inclusions that are immunoreactive against both TDP-43 and ANXA11 have been reported in these cases^33,36,37^. This suggests that pathogenic *ANXA11* variants might promote the co-assembly of ANXA11 and TDP-43 in heteromeric amyloid filaments. Individuals carrying pathogenic variants in genes encoding other LCD-containing proteins, such as hnRNPA1/A2B1 and ataxin-2 (refs. ^38,39^), present with brain inclusions that are immunoreactive against both TDP-43 and the mutated protein. This raises the possibility that TDP-43 may form heteromeric amyloid filaments with these proteins in such cases. The individuals studied here had wild-type *ANXA11*. This suggests that, in addition to FTLD-TDP Type C, subtypes of ALS and inclusion body myopathy characterised by heteromeric amyloid filaments of wild-type ANXA11 and TDP-43 may exist. This work should motivate histological examination of additional TDP-43 diseases for ANXA11 pathology.

The discovery of a second protein in the filaments of FTLD-TDP Type C offers new avenues to investigate of the enigmatic mechanisms of protein assembly in neurodegenerative diseases. The extensive hydrophobic interface between ANXA11 and TDP-43 at the centre of the filament fold strongly suggests that the two proteins co-assemble, rather than forming individual protofilaments. Co-assembly is supported by the absence of homotypic filaments of ANXA11 or TDP-43. ANXA11 and TDP-43 co-exist in axonal RNP granules under physiological conditions^40,41^, suggesting that this is where the two proteins might co-assemble. This may explain the distinct distribution of the filaments within neuritic inclusions in the prefrontal cortex in FTLD-TDP Type C.

The LCD of ANXA11 is required for its association with RNP granules^41^, but the underlying interactions are unknown. Our discovery of a pathological interaction between the LCDs of ANXA11 and TDP-43 raises the question of whether this represents the dysregulation of a physiological interaction. The regions of ANXA11 and TDP-43 that interact in the filament fold both form amphipathic α-helices in solution and have been implicated in protein-protein interactions^42,31,26,43–45^. Possibly, such interactions might precede amyloid co-assembly. Promoting interactions with other binding partners and increasing helical propensity may represent a strategy to prevent co-assembly. Future work should focus on producing model systems that recapitulate the heteromeric amyloid filament structure of ANXA11 and TDP-43 in order to investigate these hypotheses.

We found that ANXA11 predominantly exists as a previously undescribed NTF in the heteromeric filaments. Its mass of ∼22 kDa, together with the presence of residues L39–L74 in the filament fold, indicate that the NTF lacks the phospholipid-binding annexin core domain. Removal of this domain might facilitate co-assembly with TDP-43 by producing a pool of non-membrane-associated ANXA11. Alternatively, truncation might occur as a response to filament formation, as has been suggested for TDP-43^46^. Future studies should focus on the molecular mechanisms of ANXA11 truncation. The ANXA11 NTF may also facilitate the search for biomarkers for FTLD-TDP Type C, which are currently lacking.

Heteromeric interactions may also be relevant for TDP-43 assembly in other diseases, as well as for the assembly of other disease-associated proteins. Distinct filament folds are found in different diseases^1^, but the mechanisms of their formation are unknown. The interface between ANXA11 and TDP-43 in FTLD-TDP Type C filaments is incompatible with the TDP-43 folds of ALS and FTLD-TDP Type A and B^3,4^, suggesting that heteromeric interactions may be one way to influence folds.

Isolated peptides associated with the filament folds of homomeric amyloid filaments from patient brain have been observed in cryo-EM structures^4,47^. Their short lengths precluded their identification. They may derive from the same proteins that make up the filament folds, similar to the isolated peptide associated with TDP-43 residues F289–R293 described here. Alternatively, our findings raise the possibility that these peptides may be derived from other proteins. Their identification may have similar implications for our understanding of assembly mechanisms in disease.

The heteromeric amyloid filament structure of ANXA11 and TDP-43 explains the co-localisation of these two proteins in the inclusions of FTLD-TDP Type C. However, it is important to make the distinction that co-localisation of a given protein with inclusions is not sufficient evidence for heteromeric assembly, since many additional non-assembled proteins are sequestered in inclusions^48–50^, and all previously determined structures of protein assemblies in neurodegenerative diseases are homotypic amyloid filaments^1,30^. Cryo-EM is

currently unique in its ability to conclusively demonstrate that a given protein assembles in disease.

## Conclusion

The co-assembly of ANXA11 and TDP-43 in heteromeric amyloid filaments in FTLD-TDP Type C revises our understanding of amyloids as self-assemblies of identical or near-identical sequences. The relevance of this finding to other neurodegenerative diseases needs to be examined. This work establishes a central role for ANXA11 in FTLD-TDP Type C. Targeting the co-assembly of ANXA11 and TDP-43 may represent a selective and specific strategy for the diagnosis and treatment of this disease.

## METHODS

### Human tissue samples

We studied fresh-frozen, post-mortem brain tissue from nine neuropathologically-confirmed cases of FTLD-TDP Type C, three cases of FTLD-TDP Type A and three cases of FTLD-TDP Type B. Neuropathological diagnosis of FTLD-TDP Type was made according to the criteria set out in^11^. The clinicopathological details of the four FTLD-TDP Type C cases used for cryo-EM are shown in Extended Data Table 1. All FTLD-TDP Type C cases had wild-type *ANXA11* and clinical presentations of svPPA. All FTLD-TDP Type A cases carried *GRN* variants associated with FTLD-TDP Type A and had clinical presentations of behavioural variant FTD (bvFTD). Two of the FTLD-TDP Type B cases carried hexanucleotide repeat expansions in *C9orf72* and had clinical presentations of bvFTD. The third FTLD-TDP Type B case had wild-type *C9orf72* and a clinical presentation of ALS and bvFTD. The use of the tissue samples in this study was approved by the ethical review processes at each institution. Informed consent was obtained from the patients’ next of kin.

### Genetic sequencing

Whole-exome sequencing target enrichment used the SureSelectTX human all-exon library (V6, 58 megabase pairs; Agilent) and high-throughput sequencing was carried out using a HiSeq 4,000 (sx75 base-pair paired-end configuration; Illumina). Repeat-primed PCR followed by fragment length analyses were performed to screen for hexanucleotide repeat expansions within the *C9orf72* gene, as described^51^. Oligonucleotides designed to amplify the coding exons and corresponding flanking intronic regions of the *ANXA11* gene were used for polymerase chain reactions using 50 ng of genomic DNA extracted from brain tissue. The amplified products were purified and underwent direct dideoxy sequencing as described^52^.

### Amyloid filament extraction

Extraction of amyloid filaments from fresh frozen prefrontal and temporal cortex was performed as previously described^3^. Grey matter was dissected and homogenized using a Polytron (Kinematica) in 40 volumes (v/w) extraction buffer containing 10 mM Tris-HCl pH 7.5, 0.8 M NaCl, 10 % sucrose and 1 mM EGTA. A 25% solution of Sarkosyl in water was added to the homogenates, resulting in a final concentration of 2% Sarkosyl. Samples were incubated for 1 h at 37 °C with orbital shaking at 200 rpm. Following incubation, samples were centrifuged at 27,000 *g* for 10 min. The resulting supernatants were centrifuged at 166,000 *g* for 20 min. Supernatants were discarded and pellets containing TDP-43 filaments were resuspended in 6 mL/g tissue of extraction buffer containing 1% Sarkosyl by sonication for 5 min at 50% amplitude (Qsonica Q700), followed by 4-fold dilution with the same buffer and incubation for 30 min at 37 °C with orbital shaking at 200 rpm. Samples were then centrifuged at 17,000 *g* for 5 min and pellets discarded. The supernatants were further centrifuged at 166,000 *g* for 20 min followed by resuspension in 1 mL/g tissue of extraction buffer containing 1% Sarkosyl by incubation for 1 h at 37 °C with orbital shaking at 200 rpm. The samples were centrifuged at 166,000 *g* for 20 min and the final pellets were resuspended in 30 μL/g tissue of 20 mM Tris-HCl pH 7.4, 150 mM NaCl by sonication for 5 min at 50% amplitude (Qsonica Q700). 1–2 g of tissue was used for each cryo-EM sample. All centrifugation steps were carried out at 25 °C.

### Immunolabelling

For immunohistochemistry, brain hemispheres were fixed with 10 % buffered formalin, sectioned and embedded in paraffin. 8 μm-thick deparaffinized sections were incubated in 10 mM sodium citrate buffer at 105 °C for 10 min and treated with 95 % formic acid for 5 min. Sections were washed and blocked with 10 % fetal calf serum (FCS) in PBS. For colourimetric immunohistochemistry, sections were incubated overnight with a primary antibody against pS409/S410 TDP-43 (CosmoBio CAC-TIP-PTD-M01A, 1:1,000) in blocking buffer. Sections were then washed and incubated with biotinylated secondary antibodies for 2 h. Labelling was detected using an ABC staining kit (Vector) with DAB. Sections were counterstained with haematoxylin. For multiplexed fluorescent immunohistochemistry, sections were incubated overnight with primary antibodies against pS409/410 TDP-43 (CosmoBio CAC-TIP-PTD-M01A, 1:200 or 1:1,000) and N-terminal Annexin A11 (residues 1–180, Proteintech, 10479-2-AP, 1:200 or 1:1,000) in blocking buffer. Sections were washed and fluorescent secondary antibodies conjugated to Alexa Fluor 488, 568 or 594 (Thermo A11008, A32723, A11004 and A32740) were added for 2 h, followed by washing. Sections were then treated with 0.1% Sudan Black B (Fujifilm Wako) for 10 min and mounted with Vectashield with DAPI (Vector Laboratories) or ProLong Gold with DAPI (Thermo).

For immunoblotting, filaments extracted from equal amounts of initial grey matter were disassembled and denatured by incubation with LDS sample buffer containing 4% β-mercaptoethanol for 5 minutes at 95 °C. Samples were then resolved using 12% Bis-Tris gels (Novex) at 200 V for 45 min and transferred onto nitrocellulose membranes. Membranes were blocked in PBS containing 1% BSA and 0.2% Tween for 30 min at 21°C and incubated with primary antibodies against pS409/410 TDP-43 (CosmoBio CAC-TIP-PTD-M01A, 1:3,000) and N-terminal Annexin A11 (residues 1–180, Proteintech, 10479-2-AP, 1:1,000) at 21°C for 1 h. Membranes were then washed three times with PBS containing 0.2% Tween, 5 min each wash, and incubated with fluorescent secondary antibodies conjugated to StarBright Blue 520 (Bio-Rad) or DyLight 800 (Cell Signalling Technologies). Membranes were then washed three times as above and imaged using a ChemiDoc MP (Bio-Rad).

For immuno-EM, filament extracts were deposited onto carbon-coated 300-mesh copper or nickel grids (Nissin EM and Electron Microscopy Sciences, respectively), blocked with 0.1 % gelatin in PBS, and incubated with primary antibodies against pS409/410 TDP-43 (CosmoBio CAC-TIP-PTD-M01A, 1:50) and N-terminal Annexin A11 (Proteintech, 10479-2-AP, 1:50) in PBS containing 0.1% gelatin at 21 °C for 3 h or at 4°C overnight. After washing with 0.1% gelatin in PBS, the grids were incubated with secondary antibodies conjugated to 10 nm or 6 nm gold particles (Cytodiagnostics, 1:20, and Electron Microscopy Sciences, 1:40) in PBS containing 0.1% gelatin at 21 °C for 1 h. The grids were then stained with 2% uranyl acetate or NanoVan (Ted Pella). Images were acquired using 80 keV JEOL JEM-1400 and FEI Tecnai Spirit Bio-Twin electron microscopes equipped with CCD cameras.

### Mass spectrometry

Pelleted filaments extracted from 0.1 g of tissue were disassembled by resuspension in 100 μl hexafluoroisopropanol and inclubation overnight at 37°C with shaking at 200 rpm. The samples were then sonicated for 5 min at 50% amplitude (QSonica Q700) and centrifuged at 166,000 *g* for 15 min. The supernatant containing disassembled filaments was collected and dried by vacuum centrifugation (Savant). A solution of 8 M urea in 50 mM ammonium bicarbonate was added to the dried protein samples and then reduced with 5 mM DTT at 56 °C for 30 min and alkylated with 10 mM iodoacetamide in the dark at room temperature for 30 min. Samples were diluted to 1 M urea with 50mM ammonium bicarbonate and digested with chymotrypsin (Promega) at 25°C overnight. The chymotrypsin was inactivated with formic acid (FA), to a final concentration of 0.5%. Samples were then centrifuged at 16,000 *g* for 5 min.

The resulting supernatants were desalted and fractionated using custom-made C18 stop-and-go-extraction (STAGE) tips (3M Empore) packed with porous oligo R3 resin (Thermo Scientific). STAGE tips were equilibrated with 80% acetonitrile (MeCN) containing 0.5% FA, followed by 0.5% FA. Bound peptides were eluted stepwise with increasing MeCN concentrations from 5-60% MeCN in 10 mM ammonium bicarbonate, and partially dried down by vacuum centrifugation (Savant).

Fractionated peptides were analysed by LC-MS/MS using a fully automated Ultimate 3000 RSLC nano System (Thermo Scientific). Peptides were trapped with a PepMap100 C18 5 μm 0.3X5 mm nano trap column (Thermo Fisher Scientific) and an Aurora Ultimate TS 75μmx25cmx1.7μm C18 column (IonOpticks) using a binary gradient consisting of 0.1% FA (buffer A) and 80% MeCN in 0.1% FA (buffer B) at a flow rate of 300 nl/min. Eluted peptides were introduced directly via a nanoFlex ion source into an a Q Exactive Plus hybrid quadrupole-Orbitrap mass spectrometer (Thermo Scientific). MS1 spectra were acquired at a resolution of 70K; mass range of 380-1500 m/z; AGC target of 1e6; MaxIT of 100 ms; this was followed by MS2 acquisitions of the 15 most intense ions with a resolution of 17.5K. NCE of 27% and isolation window =1.2 m/z were used. Dynamic exclusion was set for 30s.

LC-MS/MS data were searched against the human reviewed database (UniProt, downloaded 2019) using the Mascot search engine (Matrix Science, v2.4). Database search parameters were set with a precursor tolerance of 10 ppm and a fragment ion mass tolerance of 0.1 Da. A maximum of three missed chymotrypsin cleavages were allowed. Carbamidomethyl cysteine was set as static modification. Arginine citrullination and methylation; and asparagine and glutamine deamination were specified as variable modifications. Scaffold (version 4, Proteome Software Inc.) was used to validate MS/MS-based peptide and protein identifications. MS/MS spectra containing arginine citrullination and methylation were manually confirmed using the Scaffold fragmentation tables.

### Cryo-EM

Filament extracts were incubated with 0.4 mg/mL pronase (Sigma) for 1 h at 21 °C. Samples were centrifuged at 3,000 *g* for 15 s and supernatants were retained. 3 μl of sample was applied to glow-discharged 1.2/1.3 μm holey carbon-coated 300-mesh gold grids (Quantifoil) and plunge-frozen in liquid ethane using a Vitrobot Mark IV (Thermo Fisher). Images were acquired using a 300 keV Titan Krios microscope (Thermo Fisher) equipped with a K3 detector (Gatan) and a GIF-quantum energy filter (Gatan) operated at a slit width of 20 eV. Aberration-free image shift (AFIS) within the EPU software (Thermo Fisher) was used during image acquisition. Further details are given in Extended Data Table 2.

### Helical reconstruction

Movie frames were gain-corrected, aligned, dose-weighted and summed using the motion correction program in RELION-4.0 or RELION-5.0^53^. The motion-corrected micrographs were used to estimate the contrast transfer function (CTF) using CTFFIND-4.1^54^. All subsequent image-processing was performed using helical reconstruction methods in RELION-4.0 or RELION-5.0^55,56^. Filaments were picked manually. Reference-free two-dimensional (2D) classification was performed to remove suboptimal segments. Initial three-dimensional (3D) reference models were generated *de novo* by producing sinograms from 2D class averages as previously described^57^. 3D auto-refinements with optimization of the helical twist were performed, followed by Bayesian polishing and CTF refinement^53,58^. 3D classification was used to further remove suboptimal segments, as well as to separate segments with alternative conformations. 3D auto-refinement, Bayesian polishing, and CTF refinement were then repeated. The final reconstructions were sharpened using the standard post-processing procedures in RELION. The overall resolutions were estimated from Fourier shell correlations of 0.143 between the two independently refined half-maps, using phase-randomization to correct for convolution effects of a generous, soft-edged solvent mask^59^. Local resolution estimates were obtained using the same phase-randomization procedure, but with a soft spherical mask that was moved over the entire map. Helical symmetry was imposed using the RELION Helix Toolbox. Further details are given in Extended Data Table 2.

### Atomic model building and refinement

The ANXA11 chain was identified by deriving the 11-residue sequence motif G[NMQ]X[SA][EDRKQN]M[SA][SAG]X[WF][SAG] from the high-resolution cryo-EM reconstructions by inspection of well-resolved densities for amino acid side chains. This motif was then searched against the combined UniProtKB and Swiss-Prot reference proteomes. The search returned ANXA11 as the single hit and enabled us to build residues L39–L74 of ANXA11 into the cryo-EM reconstruction *de novo*. We confirmed this result by using the automated machine-learning approach of ModelAngelo^60^ to calculate an initial atomic model without a reference sequence, which also returned residues L39–L74 of ANXA11. The atomic model of the TDP-43 chain and its alternative conformation were built *de novo*. The complete atomic models were refined in real-space in COOT^61^ using the best-resolved maps. Rebuilding using molecular dynamics was carried out in ISOLDE^62^. The models were refined in Fourier-space using REFMAC5^63^, with appropriate symmetry constraints defined using Servalcat^64^. To confirm the absence of overfitting, the model was shaken, refined in Fourier-space against the first half-map sing REFMAC5 and compared to the second half map. Geometry was validated using MolProbity^65^. Molecular graphics and analyses were performed in ChimeraX^66^. Model statistics are given in Extended Data Table 2.

## Data availability

Whole-exome data have been deposited in the National Institute on Ageing Alzheimer’s Disease Data Storage Site (NIAGADS) under accession code NG00107. Mass spectrometry data will be deposited to the Proteomics Identifications (PRIDE) database prior to publication. Cryo-EM datasets will be deposited to the Electron Microscopy Public Image Archive (EMPIAR) prior to publication. Cryo-EM reonstructions have been deposited to the Electron Microscopy Data Bank (EMDB) under accession code EMD-50628 and EMD-50621 (Individual 1, alternative conformations 1 and 2, respectively). Atomic models have been deposited to the Protein Data Bank (PDB) under accession codes 9FOR and 9FOF (alternative conformations 1 and 2, respectively).

## ACKNOWLEDGEMENTS

We thank the individuals and their families for donating brain tissue; the Brain Library of the Dementia Laboratory at Indiana University School of Medicine for supplying tissue from FTLD-TDP Type C individuals 2, 5 and 6, FTLD-TDP Type A individuals 1 and 2, and FTLD-TDP Type B individuals 1 and 2; the Alzheimer’s Disease Research Center at the Mesulam Center for Cognitive Neurology and Alzheimer’s Disease, Feinberg School of Medicine, Northwestern University for supplying tissue from FTLD-TDP Type C individuals 7–9; the Center for Electron Microscopy (iCEM) at Indiana University School of Medicine for support with immuno-EM; the Center for Medical Genomics of Indiana University School of Medicine for next-generation DNA sequencing; staff at the MRC Laboratory of Molecular Biology Electron Microscopy Facility for access to and support with cryo-EM; and staff at the MRC Laboratory of Molecular Biology Scientific Computing Facility for access to and support with computing; A. Bertolotti, R. Chen, S.W. Davies, A. Giblin, M. Goedert, S.H.W. Scheres, S. Tetter and N. Varghese for discussions. This work was supported by the Medical Research Council, as part of United Kingdom Research and Innovation (also known as UK Research and Innovation) (MC_UP_1201/25 to B.R.-F.); the Hans Und Ilse Breuer Stiftung (to B.R.-F.); the US National Institutes of Health (R01NS137469 to K.N., L.C. and B.R.-F; P30AG072977, R01AG077444, R01DC008552, P30AG13854, R01AG056258 and R01NS085770 to C.G. and M.M.; R01-NS110437, RF1-AG071177 and R01-AG080001 to R.V. and B.G); the Japan Agency for Medical Research and Development (AMED) (JP20dm0207072 to M.H.; JP21wm0425019 to Y.S. and S.M.); the Japan Science and Technology Agency (JST) Core Research for Evolutional Science and Technology (CREST) (JPMJCR18H3 to M.H.); the Japan Society for the Promotion of Science (JSPS) KAKENHI (JP22H04923 (CoBiA) to Y.S. and S.M.); the Integrated Research Initiative for Living Well with Dementia (IRIDE) of the Tokyo Metropolitan Institute for Geriatrics and Gerontology IRIDE (to Y.S. and S.M.); and The Leverhulme Trust (ECF-2022-610 to D.A.). For the purpose of open access, the MRC Laboratory of Molecular Biology has applied a CC BY public copyright licence to any Author Accepted Manuscript version arising.

## AUTHOR CONTRIBUTIONS

I.K., H.S., M.O., Y.S., S.M., C.G., K.L.N., M.M., B.G. and M.H identified individuals and performed neuropathological examinations. T.N., M.H.J., B.G. and M.H. performed immunohistochemistry. T.N., H.J.G. and R.V. performed genetic analyses. D.A., T.N., L.C. and M.H. extracted filaments. D.A. performed immunoblot analyses. L.C. and M.H. performed immuno-EM analyses. S.Y.P.-C performed mass spectrometry analyses. D.A. collected cryo-EM data. D.A., A.G.M. and B.R.-F. analysed cryo-EM data. B.R.-F. supervised the study. All authors contributed to writing the manuscript.

## MATERIALS & CORRESPONDENCE

Correspondence and material requests should be addressed to Benjamin Ryskeldi-Falcon at.

---

**Extended Data Fig. 1:**
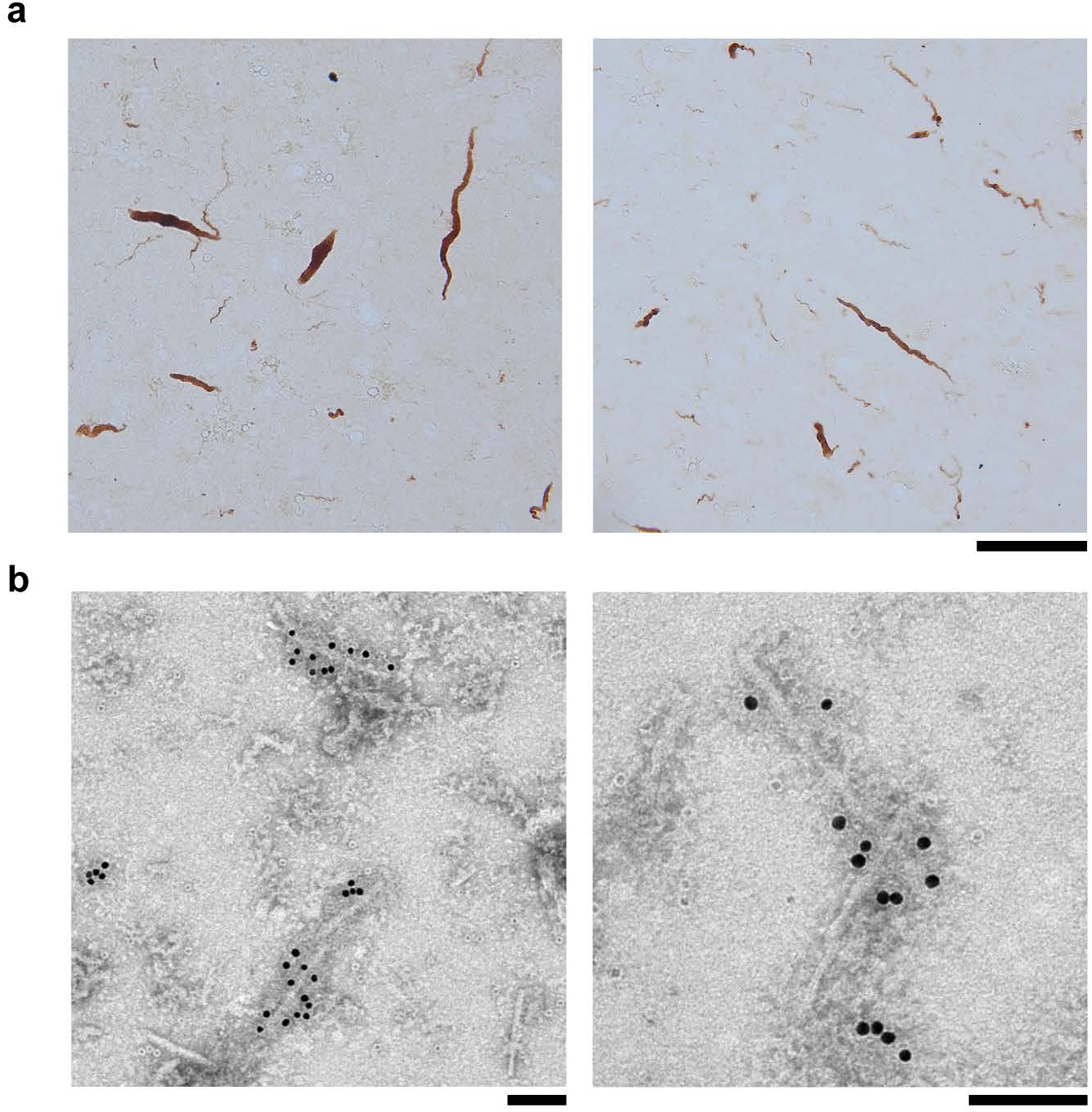
lmmunohistochemical and immuno-EM analyses of assembled TDP-43 in FTLD-TDP Type C. **a**, lmmunohistochemical analysis of prefrontal cortex sections from an individual with FTLD-TDP Type C (individual 1) using an antibody against pS409/410 TDP-43 (brown). Scale bar, 50 µm. **b,** lmmuno-EM analysis of filament extracts from the prefrontal cortex of an individual with FTLD-TDP Type C (individual 1) using an antibody against pS409/410 TDP-43 and a 10 nm gold-conjugated secondary antibody (black dots). Scale bars, 100 nm.

**Extended Data Fig. 2:**
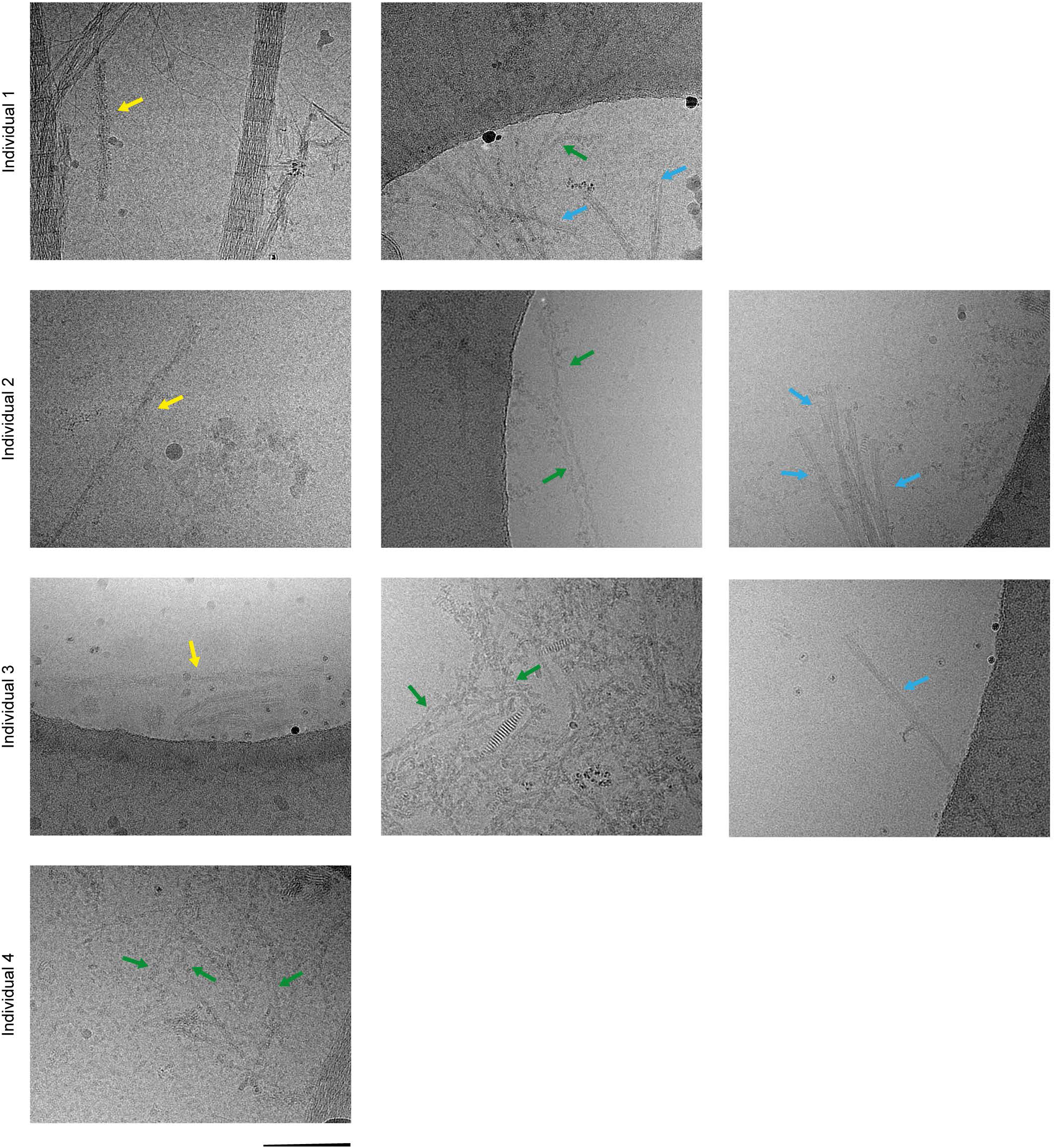
Cryo-EM of additional filament types from individuals with FTLD-TDP Type C. Representative cryo-EM images of tau paired helical filaments (yellow arrows), A filaments (green arrows) and TMEM106B filaments (cyan arrows) in the filament extracts from the prefrontal and temporal cortex of four individuals with FTLD-TDP Type C. Tau paired helical filaments were identified by their width of ∼20 nm and helical crossover distance of ∼80 nm; A filaments were identified by their width of ∼8 nm and helical crossover distance of ∼30 nm; and TMEM106B filaments were identified by their widths of ∼12 nm (single protofilament) and ∼26 nm (double protofilament), helical crossover distances of ∼200 nm and smooth surfaces. Scale bar, 100 nm.

**Extended Data Fig. 3:**
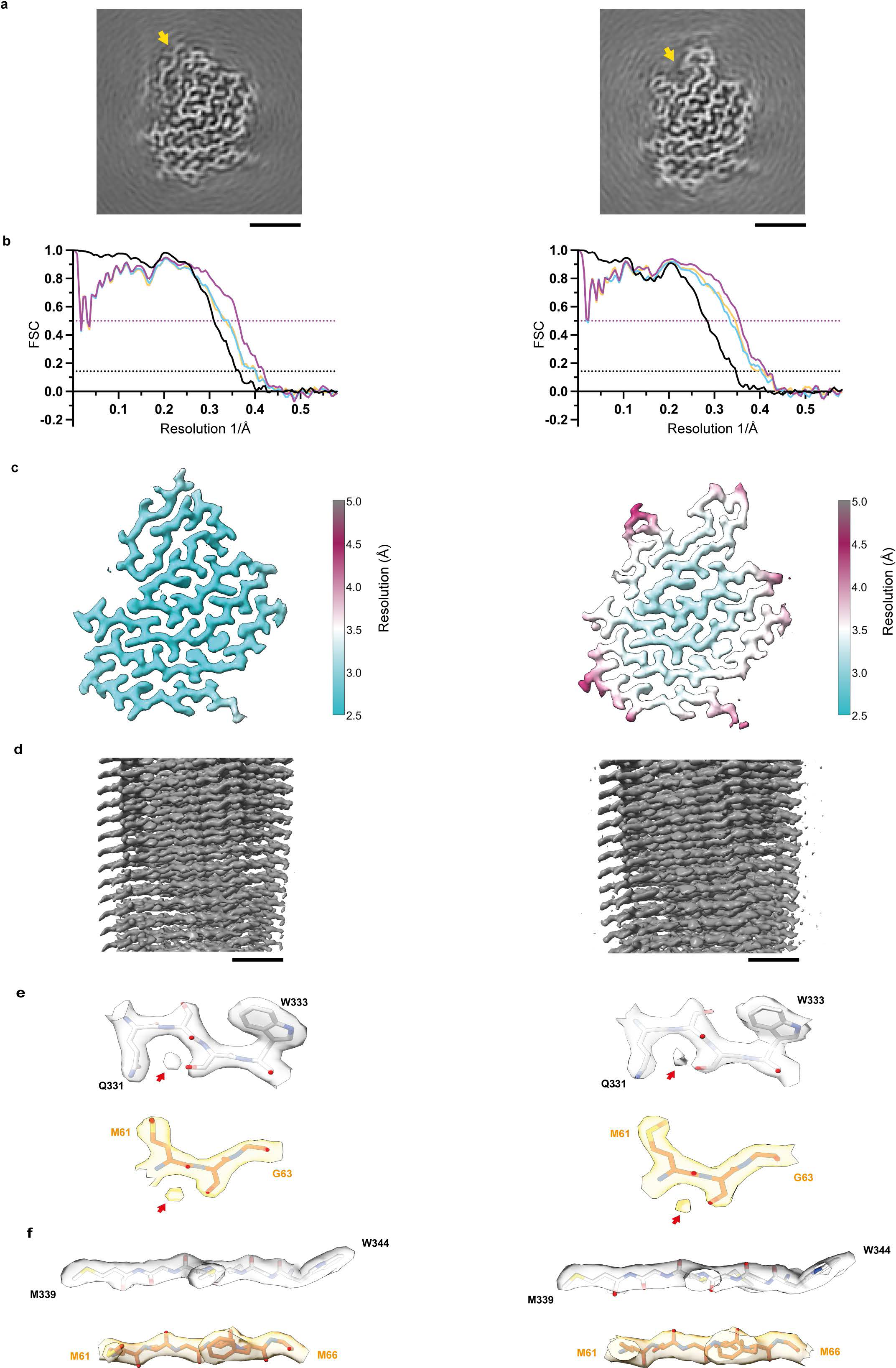
Cryo-EM reconstructions and atomic models. **a**, Cryo-EM reconstructions of filaments from **FTLD-TDP** Type C individual 1 with two alternative conformations of the TDP-43 glycine-rich region (indicated with arrows), shown as central slices perpendicular to the helical axis. Scale bars, 2 nm. **b,** Fourier shell correlation (FSC) curves for the two independently-refined cryo-EM half-maps (black lines); for the refined atomic model against the cryo-EM density map (magenta); for the atomic model shaken and refined using the first half-map against the first half-map (cyan); and for the same atomic model against the second half-map (yellow). FSC thresholds of 0.143 (black dashed line) and 0.5 (magenta dashed line) are shown. **c,** Local resolution estimates for the cryo-EM reconstructions. **d,** Cryo-EM reconstructions viewed along the helical axis. Scale bar, 1 nm. **e,f,** Views of the cryo-EM reconstructions and atomic models showing representative densities for ordered solvent (red arrows) **(f)** and main chain oxygen atoms in 13-strands, which reveal the chirality of the map.

**Extended Data Fig. 4:**
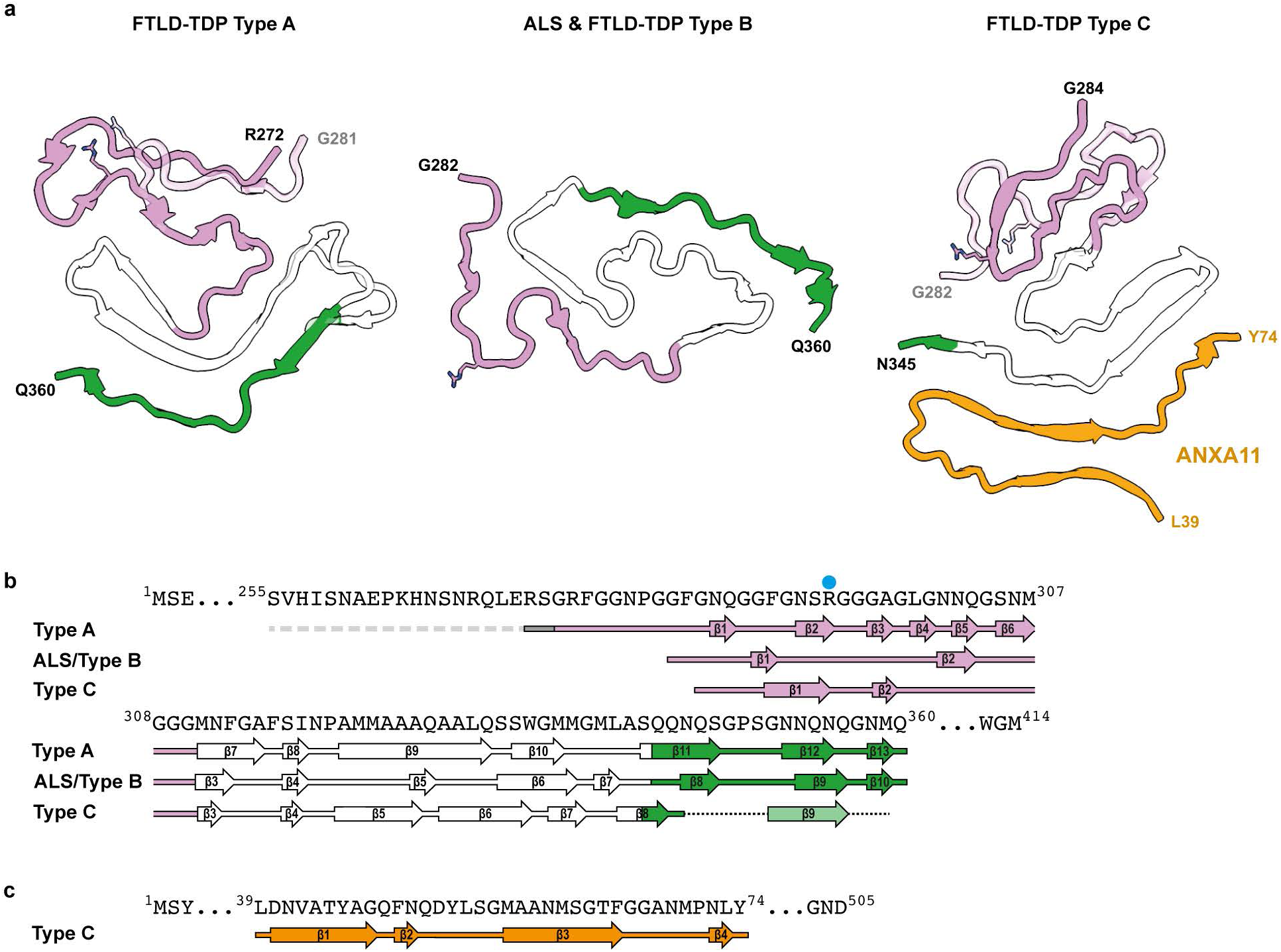
Comparison of filament folds in ALS and FTLD-TDP Type A-C. **a**, Schematic of the secondary structure elements of the homotypic TDP-43 filament folds of ALS and FTLD-TDP Type A and B, and the heterotypic TDP-43 and ANXA11 fold of FTLD-TDP Type C. Side chains for R293 are shown. Alternative local conformations of the FTLD-TDP Type A and C folds are transparent. **b and c,** Amino acid sequence alignment of the secondary structure elements of TDP-43 **(b)** and ANXA11 **(c)** in the filament folds. Arrows indicate -strands. **a-c,** The TDP-43 glycine-rich (G282-G310, magenta), hydrophobic (M311-S342, white) and QIN-rich (Q343-Q360, green) regions are highlighted. ANXA11 is shown in orange. R293 is indicated with a blue dot.

**Extended Data Fig. 5:**
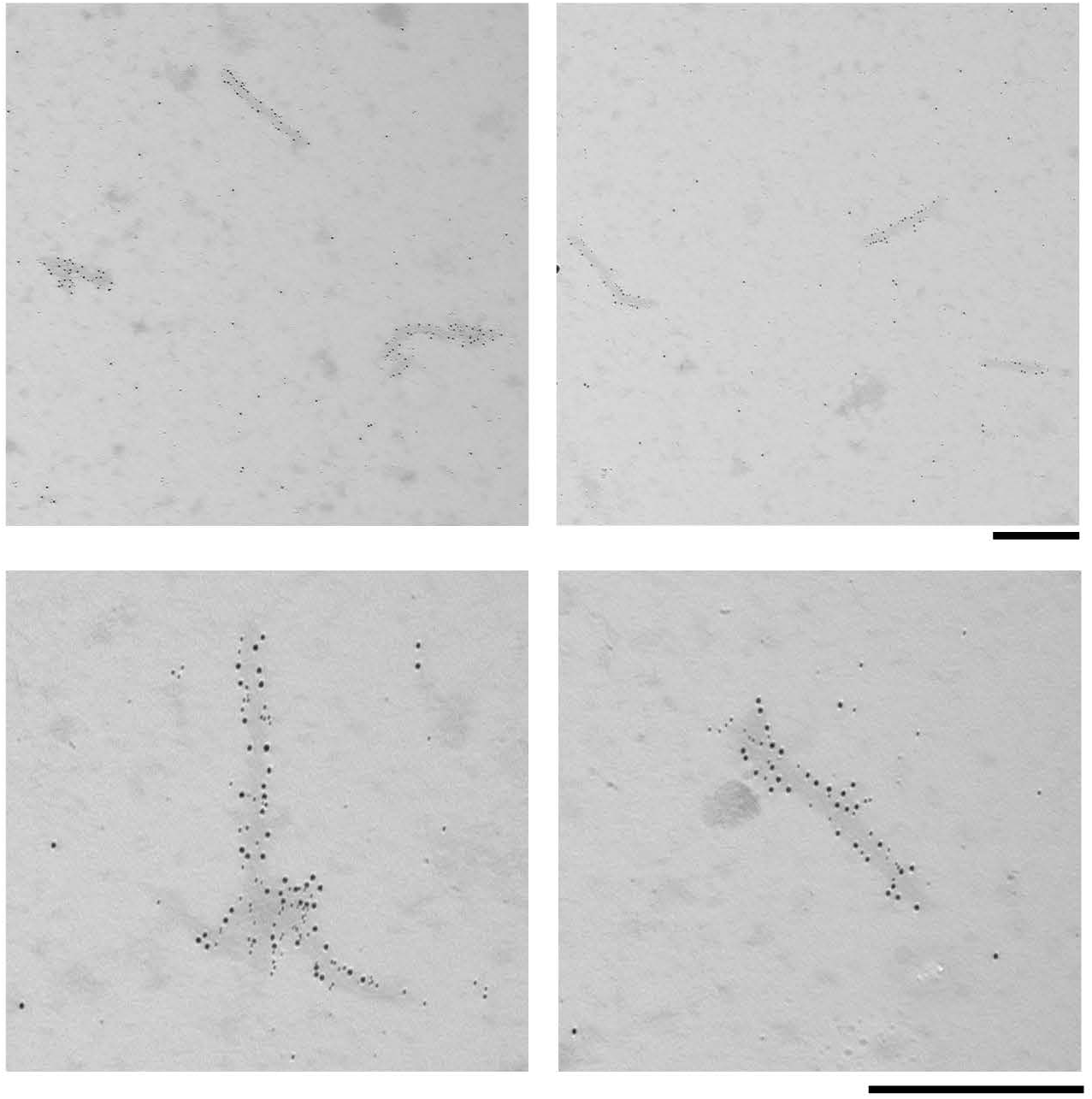
Double-labelling immuno-EM of ANXA11 and TDP-43 in FTLD-TDP Type C filaments. Double labelling immuno-EM analysis of filament extracts from the prefrontal cortex of an individual with FTLD-TDP Type C (individual 2) using antibodies against pS409/410 TDP-43 and N-terminal ANXA11 (residues 1-180) using 10 nm and 6 nm gold-conjugated secondary antibodies (black), respectively. The filaments label for both ANXA11 and TDP-43. Scale bars, 500 nm.

**Extended Data Fig. 6:**
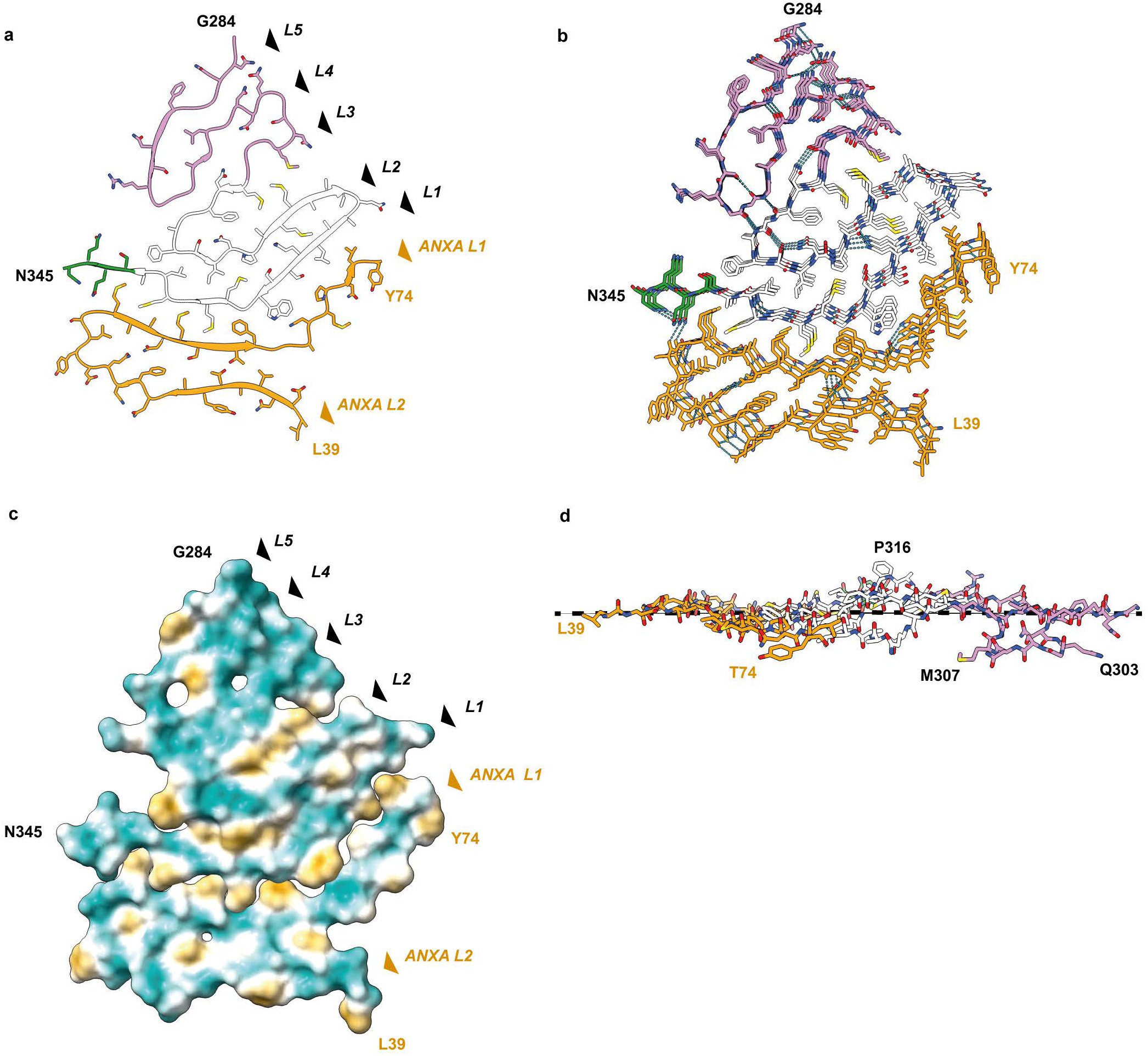
The heteromeric filament fold of ANXA11 and TDP-43 from FTLD-TDP Type C. **a**, Secondary structure of the heteromeric filament fold of FTLD-TDP Type C, shown for single ANXA11 and TDP-43 molecules perpendicular to the helical axis. **b,** Atomic model of the filament fold depicting hydrogen bonding (dashed cyan lines), shown for three ANXA11 and TDP-43 molecules perpendicular to the helical axis. **c,** Hydrophobicity of the filament fold, from most hydrophilic (teal) to most hydrophobic (yellow), shown for single ANXA11 and TDP-43 molecules perpendicular to the helical axis. **d,** Atomic model of filament fold, shown for single ANXA11 and TDP-43 molecules aligned with the helical axis. **a, b and d,** The TDP-43 glycine-rich (G284-G310, magenta), hydrophobic (M311-S342, white) and QIN-rich (Q343-Q345, green) regions are highlighted. ANXA11 is shown in orange. **a and c,** The layers of the ANXA and TDP-43 chains are indicated with arrows.

**Extended Data Fig. 7:**
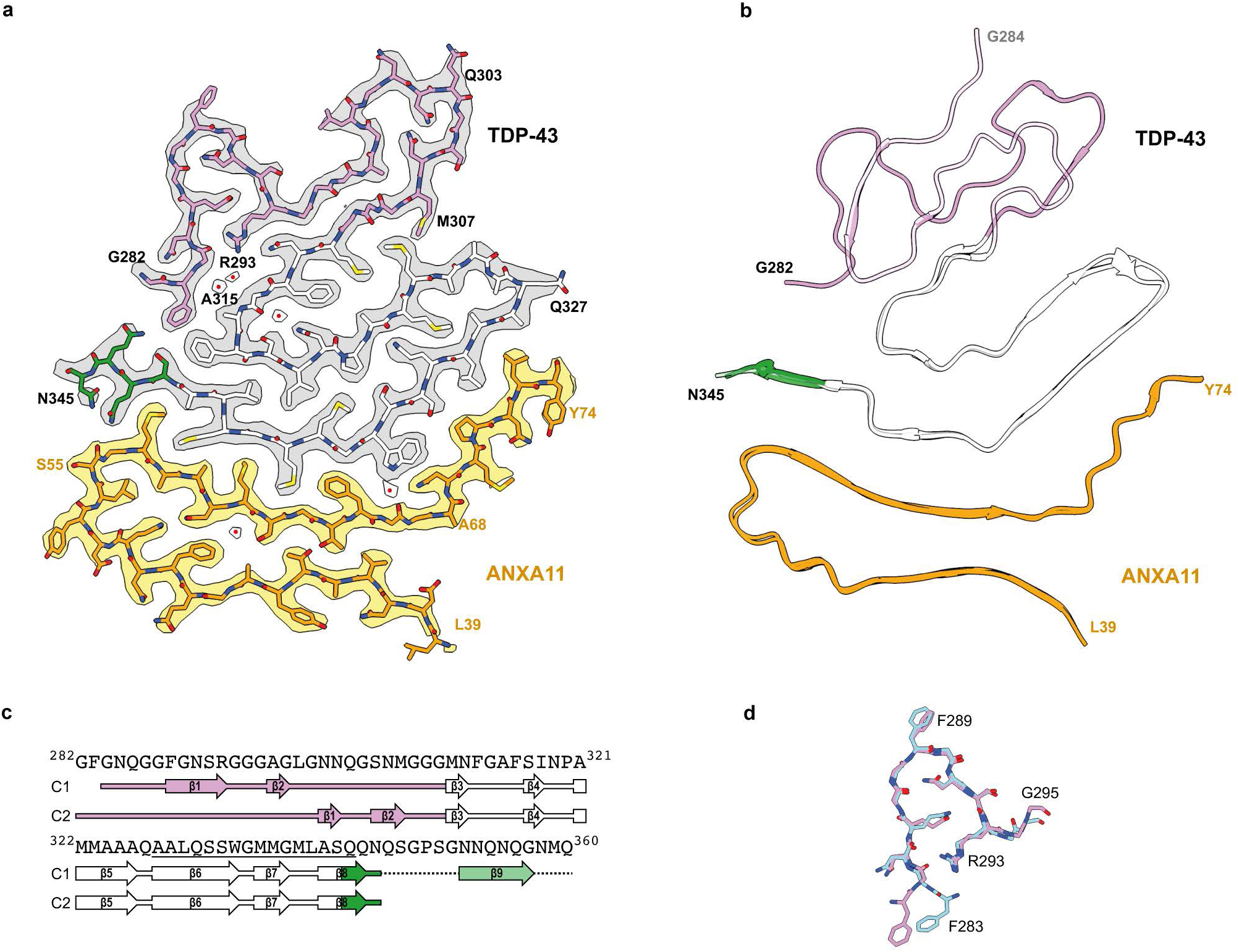
Alternative confirmation of the TDP-43 glycine-rich region in heteromeric amyloid filaments from FTLD-TDP Type C. **a**, Cryo-EM reconstruction and atomic model of heteromeric amyloid filaments from FTLD-TDP Type C with an alternative conformation of the glycine-rich region, shown for single ANXA11 and TDP-43 molecules perpendicular to the helical axis. Cryo-EM density for TDP-43 is in grey and ANXA11 is in yellow. Buried ordered solvent is indicated with red dots. **b,** Overlay of the atomic models of filaments with the alternative conformation of the glycine-rich region with the main conformation (transparent). **c,** Amino acid sequence alignment of the secondary structure elements of TDP-43 in the filaments. Arrows indicate (3-strands. C1, main conformation; C2, alternative conformation. **d,** Alignment of TDP-43 residues N295-R293 from the atomic models of FTLD-TDP Type C (pink) and Type A (cyan) filaments. **a-c,** The TDP-43 glycine-rich (G282-G310, magenta), hydrophobic (M311-S342, white) and QIN-rich (Q343-Q345, green) regions are highlighted. ANXA11 is show in orange.

**Extended Data Fig. 8:**
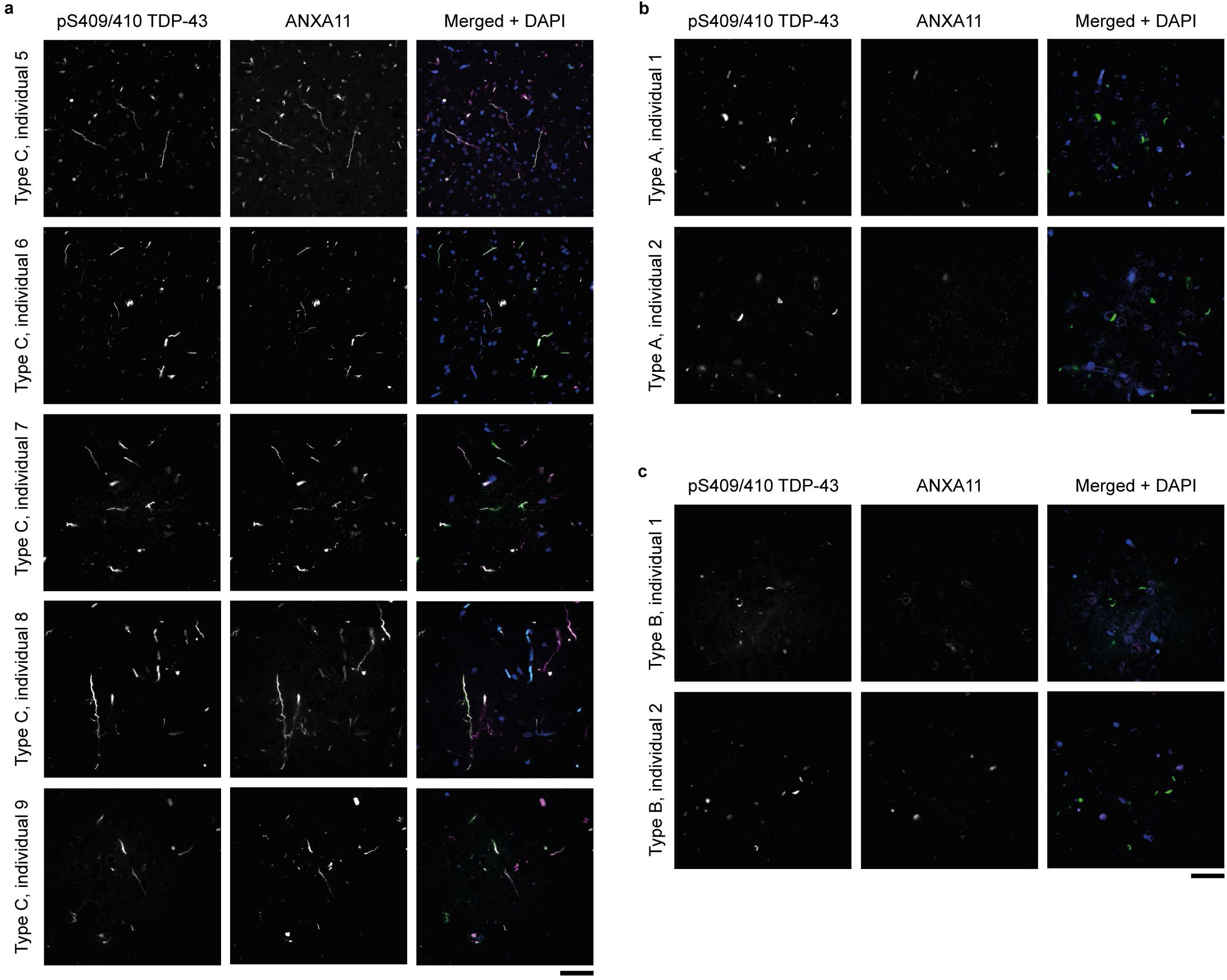
lmmunohistochemical analysis of ANXA11 and TDP-43 in the prefrontal cortex in FTLD-TDP. **a-c**, lmmunohistochemical analysis of prefrontal cortex sections from individuals with FTLD-TDP Type C **(a),** Type A **(b)** and Type B (c) using antibodies against pS409/410 TDP-43 and N-terminal ANXA11 (residues 1-180). Individual images for TDP-43 and ANXA11 are shown in greyscale to facilitate comparison, in addition to a merged image showing TDP-43 (green), ANXA11 (magenta) and DAPI (blue) staining. ANXA11 and TDP-43 colocalise with inclusions in the individuals with FTLD-TDP Type C, but only TDP-43 co-localises with the inclusions in the individuals with FTLD-TDP Types A and B. Additional immunohistochemical analysis is shown in Fig. 4c and Extended Data Fig. 9. Scale bar, 40 µm.

**Extended Data Fig. 9:**
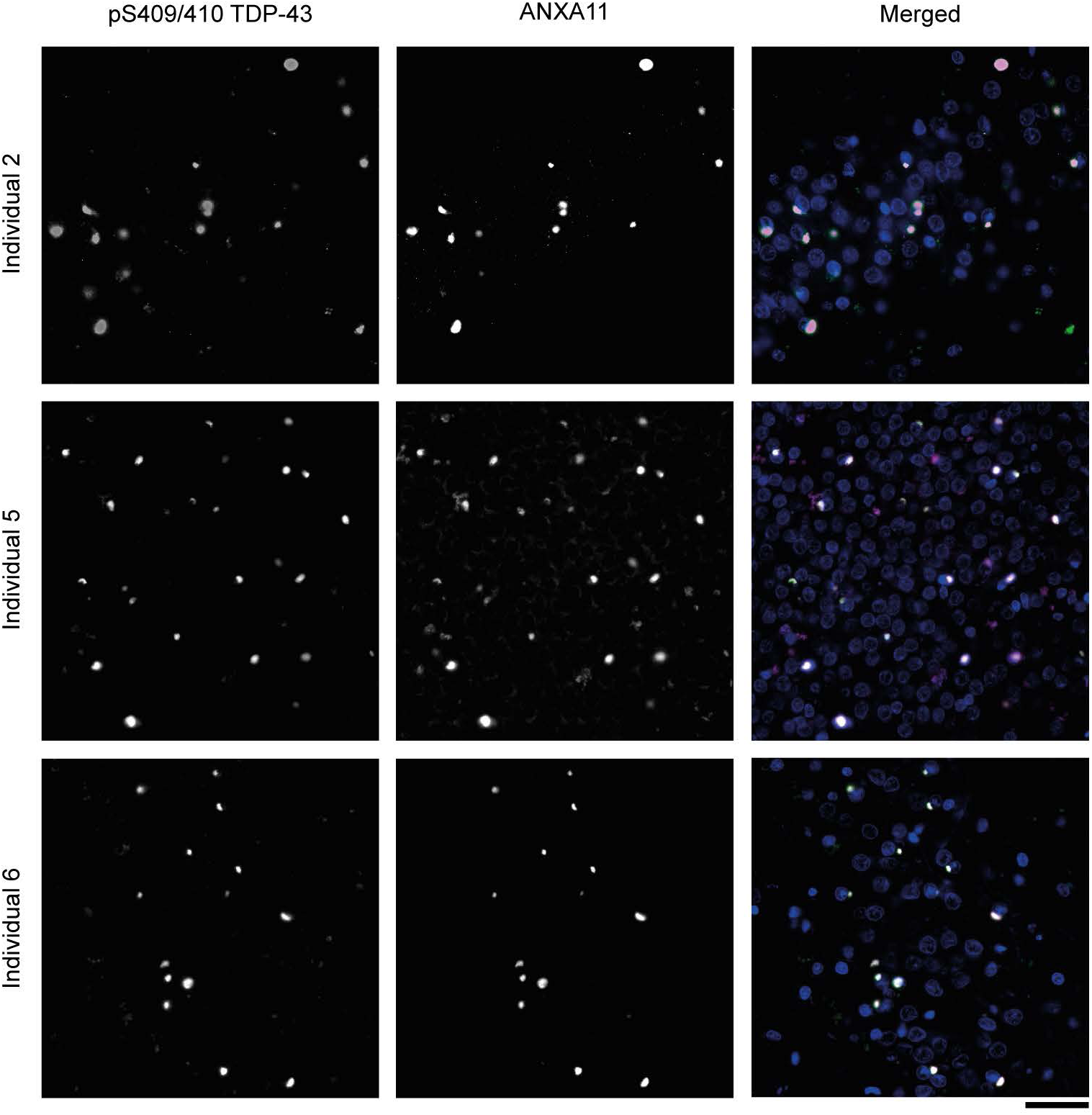
lmmunohistochemical analysis of ANXA11 and TDP-43 in the hippocampal dentate gyrus in FTLD-TDP Type C. lmmunohistochemical analysis of fascia dentata sections from three individuals with FTLD-TDP Type C using antibodies against pS409/410 TDP-43 and N-terminal ANXA11 (residues 1-180). Individual images for TDP-43 and ANXA11 are shown in greyscale to facilitate comparison, in addition to a merged image showing TDP-43 (green), ANXA11 (magenta) and DAPI (blue) staining. ANXA11 and TDP-43 colocalise with inclusions. Additional immunohistochemical analysis is shown in Fig. 4c and Extended Data Fig. 8. Scale bar, 40 µm.

**Extended Data Table 1:** Clinicopathological details.

|  | Individual 1 |  | Individual 2 |  | Individual 3 |  | Individual 4 |
| --- | --- | --- | --- | --- | --- | --- | --- |
| Male/ female | F |  | F |  | M |  | M |
| Age (y) | 81 |  | 74 |  | 86 |  | 59 |
| Disease duration (y) | 20 |  | 11.5 |  | ND |  | 12 |
| Clinical diagnosis | svPPA |  | svPPA |  | svPPA |  | svPPA |
| Neuropathological diagnosis | FTLD-TDP Type C |  | FTLD-TDP Type C |  | FTLD-TDP Type C |  | FTLD-TDP Type C |
| Brain regions studied | PFC, TC |  | PFC |  | PFC |  | PFC, TC |
| Thal phase | ND |  | 1 |  | 1 |  | ND |
| Braak stage | 0 |  | 0 |  | 1 |  | 0 |
| CERAD score | ND |  | 0 |  | 0 |  | ND |
| Lewy score | 0 |  | 0 |  | 0 |  | 0 |

F, female; M, male; y, years; svPPA, semantic variant primary progressive aphasia; FTLD-TDP, frontotemporal lobar degeneration with TDP-43 pathology; PFC, prefrontal cortex; TC, temporal cortex; ND, not determined.

**Extended Data Table 2:** Cryo-EM data collection, refinement and validation statistics.

|  | Individual 1<br>(EMPIAR-XXXXX)<br>(EMDB-50621; EMDB-50628)<br>(PDB 9FOF; PDB 9FOR) |  | Individual 2<br>(EMPIAR-XXXXX)<br>(EMDB-XXXXX) | Individual 3<br>(EMPIAR-XXXXX)<br>(EMDB-XXXXX) | Individual 4<br>(EMPIAR-XXXXX)<br>(EMDB-XXXXX) |
| --- | --- | --- | --- | --- | --- |
| Data collection |  |  |  |  |  |
| Voltage (kV) | 300 |  | 300 | 300 | 300 |
| Electron source | XFEG |  | XFEG | XFEG | XFEG |
| Detector | K3 |  | K3 | K3 | K3 |
| Electron exposure (e−/Å²) | 38 |  | 38 | 43 | 46 |
| Defocus range (µm) | -2.2 to -1.0 |  | -2.2 to -1.0 | -2.2 to -1.0 | -2.2 to -1.0 |
| Pixel size (Å) | 0.86 |  | 0.86 | 0.86 | 0.73 |
| Data processing |  |  |  |  |  |
| Initial particle images (no.) | 619,874 |  | 72,366 | 62,112 | 109,728 |
|  | Variant 1 | Variant 2 |  |  |  |
| Final particle images (no.) | 18,020 | 10,842 | 8,236 | 10,928 | 9,346 |
| Symmetry imposed | C1 | C1 | C1 | C1 | C1 |
| Helical twist (°) | -1.83 | -1.92 | -1.82 | -1.83 | -1.83 |
| Helical rise (Å) | 4.98 | 4.96 | 4.81 | 4.97 | 4.98 |
| Map resolution (Å) | 2.75 | 2.9 | 2.84 | 3.33 | 3.78 |
| FSC threshold | 0.143 |  | 0.143 | 0.143 | 0.143 |
| Map resolution range (Å) | 2.6 to 7.88 | 2.78 to 8.08 | 2.72 to 6.6 | 3.13 to 9.6 | 3.56 to 16.63 |
| Refinement |  |  |  |  |  |
| Model resolution (Å) | 2.75 | 2.9 | — | — | — |
| FSC threshold | 0.5 |  | — | — | — |
| Map sharpening *B* factor (Å²) | -47.13 | -50.183 | — | — | — |
| Model composition |  |  |  |  |  |
| Non-hydrogen atoms | 681 | 696 | — | — | — |
| Protein residues | 98 | 101 | — | — | — |
| Ligands | - |  | — | — | — |
| *B* factors (Å²) |  |  |  |  |  |
| Protein (TDP-43) | 49.5 | 47.3 | — | — | — |
| Protein (ANXA11) | 54.9 | 56.8 | — | — | — |
| Ligand | - | - |  |  |  |
| R.m.s. deviations |  |  |  |  |  |
| Bond lengths (Å) | 0.0081 | 0.0089 | — | — | — |
| Bond angles (°) | 1.974 | 1.972 | — | — | — |
| Validation |  |  |  |  |  |
| MolProbity score | 1.26 | 1.38 | — | — | — |
| Clashscore | 1.23 | 0.75 | — | — | — |
| Favoured rotamers (%) | 98.63 | 98.63 | — | — | — |
| Poor rotamers (%) | 0 | 0 |  |  |  |
| Ramachandran plot |  |  |  |  |  |
| Favored (%) | 93.62 | 93.75 | — | — | — |
| Allowed (%) | 6.38 | 5.21 | — | — | — |
| Outliers (%) | 0 | 0 | — | — | — |

